# Guarding versus self-guarding in innate immunity

**DOI:** 10.64898/2026.03.10.710826

**Authors:** Ben Ashby, Alyssa Anderson

**Affiliations:** Department of Mathematics, Simon Fraser University, Burnaby, BC, Canada

## Abstract

Hosts have evolved a variety of innate immune responses to pathogens. In many cases, hosts directly detect pathogen-associated molecular patterns (PAMPs) or pathogen effectors to trigger an immune response. However, hosts may also detect pathogens indirectly through ‘guarding’, whereby immune receptors (‘guards’) monitor the effects of pathogens (e.g., modification of target cells) rather than the pathogens themselves. Guarding poses a different evolutionary challenge for pathogens than direct recognition of PAMPs, as replication may necessitate the modification or disruption of guarded host proteins (‘guardees’). Recently, self-guarding has been discovered, in which the host target functions as both guard and guardee. Self-guarding appears to present an intractable problem for pathogens: modification of the host target may benefit replication, but also triggers an immune response. If self-guarding creates an apparently inescapable detection mechanism, why has self-guarding only recently been discovered? Here, we use mathematical models of within-host pathogen and immune dynamics to compare guarding and self-guarding architectures. We show that self-guarding leads to a more rapid immune response and faster pathogen suppression, but is also more prone to false-positive immune responses, likely imposing greater costs through autoimmunity. We therefore hypothesise that the greater potential for false-positive immune responses may limit the conditions under which self-guarding evolves.

## Introduction

Organisms across the tree of life have evolved a variety of innate defensive mechanisms to prevent or manage the impact of infectious diseases, from restriction–modification systems in bacteria (Vasu and Nagaraja 2013) to natural killer cells in mammals (Vivier et al. 2011). In many cases, the first line of innate host defence involves recognising pathogen-associated molecular patterns (PAMPs) (Fig. 1A) (Janeway 1989), but pathogens can employ various tactics to avoid detection by molecular patterns. For example, the bacterium *Helicobacter pylori* has evolved a structurally modified lipopolysaccharide that has reduced recognition by Toll-like receptor 4 (TLR4) (Tran et al. 2005), and coronaviruses encode a 2’-O-methyltransferase that modifies the viral RNA cap to mimic host mRNA and thus reduce detection (Züst et al. 2011).

**Figure 1:**
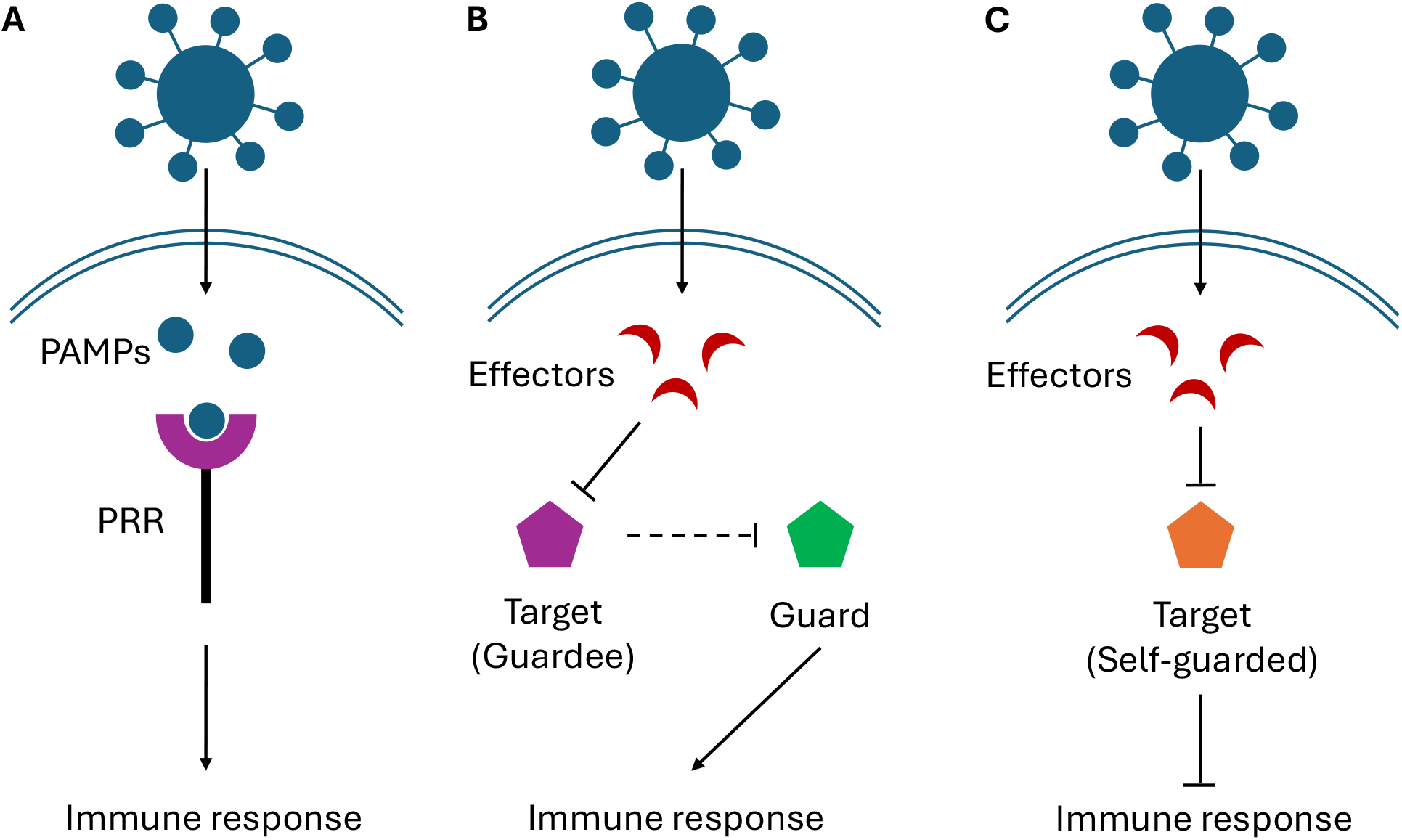
Comparison of innate immunity mechanisms. (A) Direct detection of pathogen-associated molecular patterns (PAMPs) by pathogen recognition receptors (PRRs). (B) Guarding, where pathogen effectors target a particular host protein (guardee). Another host protein (guard) monitors the guardee for perturbation (dashed arrow), which then triggers an immune response. (C) Self-guarding, where the target is a negative regulator of an immune response; modification of the self-guarded target by effectors triggers an immune response.

To combat pathogen adaptations to avoid PAMP recognition, many hosts have evolved multi-layered innate immunity targeting different aspects or stages of infection (Jones and Dangl 2006; Netea et al. 2019; Ilchenko and Pfeifer 2024). A common second layer of defence involves indirect sensing of pathogen effects on host cells or proteins rather than direct sensing of PAMPs. For example, activation of the inflammasome receptor NLRP3 can be triggered by cellular damage or ion fluxes caused by diverse microbial toxins (Lamkanfi and Dixit 2014). In plants, intracellular nucleotide-binding leucine-rich repeat (NLR) receptors monitor specific host proteins that are modified by pathogen effectors (Van der Biezen and Jones 1998; Jones and Dangl 2006). This is known as “guarding” (Dangl and Jones 2001) which often falls under the broader umbrella of effector-triggered immunity (ETI) (Remick et al. 2023).

Guarding typically occurs when particular host receptors (guards) monitor key host proteins (guardees) for modification, which leads to a secondary immune response (Fig. 1B) (Van der Biezen and Jones 1998; Dangl and Jones 2001). For example, in *Arabidopsis thaliana*, the NLR immune receptor RPS2 guards the host protein RIN4, which is cleaved by the effector AvrRpt2 during infection by *Pseudomonas syringae* (Mackey et al. 2002). However, some *P. syringae* strains have lost or modified AvrRpt2, or produce other effectors that allow them to evade guarding (Göhre and Robatzek 2008; Lindeberg et al. 2012). Vertebrates also exhibit guarding, for example via “missing self” recognition of classical MHC class I molecules by natural killer (NK) cells (Ljunggren and Kärre 1990; Lanier 2005). Many viruses downregulate classical MHC-I to evade CD8^+^ T cell responses (Yewdell and Bennink 1999; Hansen and Bouvier 2009), and this perturbation is detected by NK cell inhibitory receptors. Loss of classical MHC-I expression reduces inhibitory signalling and thereby activates NK cells (Lanier 2005). HLA-E also presents signal peptides derived from other HLA-I molecules and engages inhibitory receptors such as CD94/NKG2A (Braud et al. 1998). Reduced expression of classical HLA-I can therefore decrease HLA-E surface expression, leading to a further loss of inhibitory signalling.

Recently, novel examples of “self-guarding” have been identified, where a single protein functions as both guard and guardee (Fig. 1C) (Gaidt et al. 2021; Remick et al. 2026). The first example of self-guarding was identified in the context of herpes simplex virus type 1 (HSV-1) (Gaidt et al. 2021). HSV-1 produces the effector ICP0 to degrade the host protein MORC3, thereby relieving MORC3-mediated restriction of viral replication. But MORC3 also represses interferon (IFN) signalling. Degradation of MORC3 by ICP0 relieves negative regulation of IFN, resulting in a secondary immune response. Crucially, by having both guard and guardee functions embodied in a single protein, the virus cannot independently manipulate sensing and replication restriction functions of the host. A similar example of self-guarding against poxviruses has also been identified (Remick et al. 2026). In this case, viral targeting of antiviral host factors simultaneously disrupts their primary antiviral functions while relieving their secondary role as negative regulators of NF-*κ*B signalling. As a result, disruption of these proteins by a viral effector indirectly activates NF-*κ*B, generating an immune response that is contingent on pathogen-specific interference.

The discovery of self-guarding raises the question of when it should be favoured over separate guard and guardee functions. One possible constraint on self-guarding is that it is simply difficult or inefficient to effectively encapsulate both functions in a single protein. It is also possible that self-guarding is common but few examples have yet to be discovered. Alternatively, there may be costs inherent to self-guarding. Here, we use mathematical models of guarding and self-guarding mechanisms to explore the potential costs and benefits of these distinct architectures. We show that while self-guarding leads to faster immune responses that lead to lower peak viral loads and more rapid clearance, it is also more prone to produce false-positive immune responses, potentially leading to autoimmunity or higher metabolic costs of mounting unnecessary immune responses. In contrast, separate guard and guardee functions can reduce the risk of false-positives by smoothing out noisy signals from the guardee.

## Methods

### Model description

We model virus–host interactions at the within-host level in the presence of guarding or self-guarding. We assume that viral load (*V*) grows exponentially with per-capita growth rate *r*, but viral replication is reduced by a host regulatory factor (*R*) at pairwise rate *d*_*R*_ and by a triggered immune response (*I*) at pairwise rate *d*_*I*_. The host regulatory factor is produced at rate *s*(*R*_0_ − *R*), where *s* is the homeostatic recovery rate of *R* and *R*_0_ is its baseline (homeostatic) level. The regulatory factor is degraded or inactivated at pairwise rate *e* (e.g., by viral effectors, or due to binding with the virus to prevent replication). The immune response, *I*, is activated and deactivated at per-unit rates *m*_on_ and *m*_off_, respectively.

The specific trigger for activation of the immune response depends on the architecture of guarding. Under self-guarding, we assume that the immune response is activated at rate *m*_on_*D*(*u*(*R*)), where

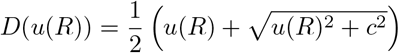

is a smooth threshold activation function which approximates the hinge function max(0, *u*(*R*)) with smoothing parameter *c*, and

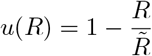

is the normalized deviation of the regulatory factor relative to the activation threshold, 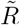. Thus, when the regulator factor falls below 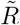, the signal induced by *D*(*u*(*R*)) transitions smoothly from zero to an approximately linear response (Fig. S1A). Under guarding, we assume that the immune response is activated at rate *m*_on_*G*, where *G* is the guard-derived activation signal. The signal *G* is produced at a rate proportional to the guard activation factor

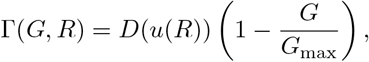

with proportionality constant *g*_on_ setting the scale of guard-signal production at rate *g*_on_Γ(*G, R*) and *G*_max_ the maximum level of the guard signal. Thus, as the regulatory factor *R* declines, the production of *G* increases, but this production saturates as *G* approaches *G*_max_. The dependence of the guard activation factor on the regulatory factor and on the saturation constant *G*_max_ is illustrated in Fig. S1B. The signal produced by the guard signal decays at rate *g*_off_. In the unsaturated (approximately linear) regime, self-guarding acts as a single first-order low-pass filter on the trigger signal, whereas guarding introduces an additional stage and thus acts as a second-order low-pass filter. The dynamics of viral load and the regulatory factor are given by

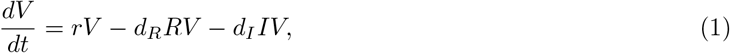

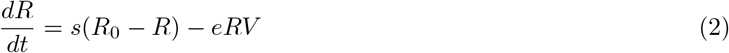

under guarding and self-guarding. The dynamics of the immune response and guarding are given by

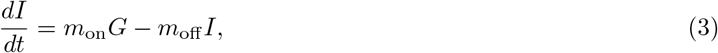

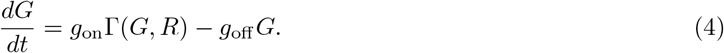

Under self-guarding, the *G* state is not present and the dynamics of the immune response are only given by

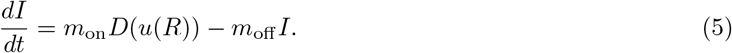

The variables and parameters for this model are summarized in Table 1. We non-dimensionalise the models by introducing the dimensionless variables and parameters described in Table 2. This yields the following non-dimensionalised equations for the dynamics of rescaled viral load (*v*) and the regulatory factor (*x*):

**Table 1:** Model variables and parameters.

| Symbol | Description |
| --- | --- |
| $V$ | Viral load |
| $R$ | Regulatory (protective) host factor |
| $I$ | Immune response |
| $G$ | Guard-generated activation signal |
| $r$ | Per-capita viral replication rate |
| $d_R$ | Pairwise suppression rate of viral replication by $R$ |
| $d_I$ | Pairwise suppression rate of viral replication by $I$ |
| $s$ | Homeostatic recovery rate of $R$ toward $R_0$ |
| $R_0$ | Baseline (homeostatic) level of $R$ |
| $\tilde{R}$ | Activation threshold level of $R$ |
| $e$ | Pairwise inactivation/degradation rate of $R$ due to interaction with $V$ |
| $m_{\text{on}}$ | Immune activation rate per unit input signal |
| $m_{\text{off}}$ | Immune deactivation rate |
| $g_{\text{on}}$ | Guard signal activation rate |
| $g_{\text{off}}$ | Guard signal deactivation rate |
| $G_{\text{max}}$ | Maximum guard signal level |
| $c$ | Immune trigger smoothing parameter |

**Table 2:** Non-dimensional variables and parameters with substitutions and dimensionless interpretations.

| Non-dimensional quantity | Interpretation (dimensionless) |
| --- | --- |
| $\tau = st$ | Time measured in units of $1/s$ |
| $x = R/R_0$ | Regulatory factor relative to baseline |
| $v = (e/s)V$ | Scaled viral load |
| $y = (d_I/r)I$ | Immune effect scaled by its impact on viral growth |
| $z = G/G_{\text{max}}$ | Guard signal relative to its maximum |
| $\rho = r/s$ | Viral replication timescale relative to $R$ recovery |
| $\alpha = d_R R_0 / r$ | Strength of regulation-mediated suppression relative to replication |
| $\theta = \tilde{R}/R_0$ | Trigger threshold relative to baseline regulation |
| $\gamma_{\text{on}} = \frac{g_{\text{on}}}{sG_{\text{max}}}$ | Guard activation rate relative to $R$ recovery and maximum guard signal level |
| $\gamma_{\text{off}} = g_{\text{off}}/s$ | Guard deactivation rate relative to $R$ recovery |
| $\lambda = \gamma_{\text{on}}/\gamma_{\text{off}}$ | Gain factor of the guard |
| $\mu_{\text{on}}^{(g)} = \frac{d_I m_{\text{on}} G_{\text{max}}}{rs}$ | Immune activation strength per unit signal (guarding) |
| $\mu_{\text{on}}^{(sg)} = \frac{d_I m_{\text{on}}}{rs}$ | Immune activation strength per unit signal (self-guarding) |
| $\mu_{\text{off}} = m_{\text{off}}/s$ | Immune deactivation rate relative to $R$ recovery |

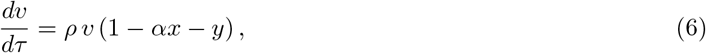

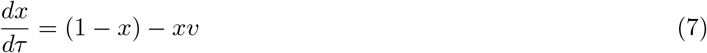

The non-dimensionalised dynamics of guarding and the immune response are then given by

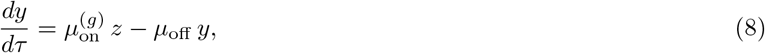

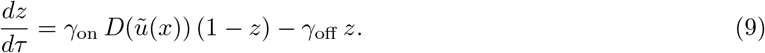

where *D*(·) is the same smooth hinge function defined above. We define *λ* = *γ*_on_*/γ*_off_ to be the ‘gain factor’ of the guard, which quantifies the relative strength of guard activation to deactivation and thus determines the extent to which the guard amplifies deviations of the regulatory factor below the activation threshold.

For self-guarding, the state *z* is not present and the dynamics of the non-dimensionalised immune response are only given by

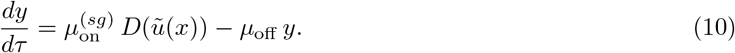

where 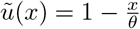, with 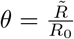 being the dimensionless threshold ratio.

#### Numerical analysis

We numerically solve the models of guarding and self-guarding using ode45 in MATLAB. Simulations are terminated when viral load falls below a nominal threshold. To explore the effects of false-positive immune activation under the guarding and self-guarding architectures, we implement a stochastic version of our model in the absence of infection (i.e., viral load is fixed at *v* = 0). This isolates spontaneous immune activation events that arise purely from intrinsic fluctuations in the regulatory factor rather than from pathogen-driven perturbations. Stochasticity is introduced only in the dynamics of the regulatory factor *x* using an additive noise term, yielding the stochastic differential equation

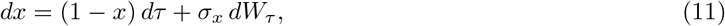

where *σ*_*x*_ controls the strength of stochastic fluctuations and *W*_*τ*_ is a standard Wiener process. This formulation corresponds to additive noise affecting perturbations away from the homeostatic level of the regulatory factor, which can represent fluctuations in transcription levels or environmental perturbations. We use additive noise for simplicity as we are only interested in relatively small deviations from homeostasis; multiplicative noise (where perturbations are proportional to regulator abundance) yields similar results. The remaining state variables for the immune response and guarding (when applicable) are treated deterministically based on the signal from *x*. The stochastic simulations are performed using an Euler–Maruyama discretisation scheme (Kloeden and Platen 1992). To ensure that *x* remains biologically meaningful, we impose hard bounds such that *x* ∈ [0, 1]. Since *v* = 0, any immune activation (*y >* 0) represents a false-positive response induced solely by stochastic excursions of *x* below its activation threshold *θ*.

## Results

We first explore analytic approximations to the dynamics of guarding and self-guarding to compare their effects on the immune response. Under self-guarding, the dynamics of the immune response are given by

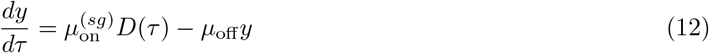

where *D*(*τ*) = *D*(*ũ*(*x*(*τ*))) is the trigger signal. Thus, under self-guarding the immune system effectively integrates the trigger signal over time with a single characteristic timescale 1*/µ*_off_. The explicit solution is then

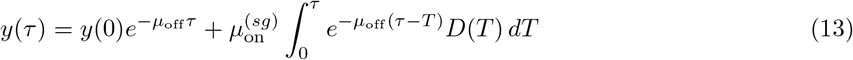

which means that the immune response is given by a convolution of the trigger signal with an exponential kernel. Self-guarding therefore acts as a first-order low-pass filter on the trigger signal *D*(*τ*). In other words, the immune response is more sensitive to slowly varying signals from *D*(*τ*) and rapid variation is attenuated. Note that the dynamics of the regulatory factor occur on a timescale of order unity due to non-dimensionalisation, and so the natural timescale of *D* is also typically 𝒪 (1). The immune response therefore has characteristic decay time

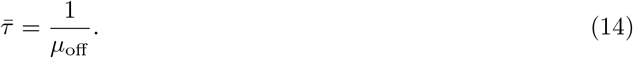

Biologically, the immune response is likely to occur on a comparable or slower timescale than regulatory dynamics, and so *µ*_off_ ≲ 1. Hence, rapid variations in the regulatory factor are attenuated, smoothing the immune response.

Under normal guarding architecture, with distinct guards and guardees, the dynamics of the guard activation signal are given by:

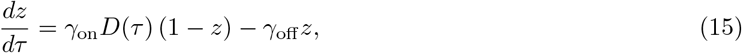

with immune activation following equation (8). The trigger signal *D*(*τ*) first drives the guard activation signal *z*, which in turn activates the immune response *y*. The guard dynamics initially follow

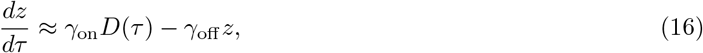

when *z* ≈ 0, which is a first-order linear filter of the trigger signal with characteristic timescale

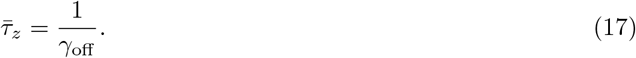

in the linear regime. The guard therefore introduces a second (intermediate) low-pass filter into the immune response provided the guard dynamics are unsaturated. This additional filtering stage has two important dynamical consequences. First, high-frequency fluctuations in *D*(*τ*) are more strongly attenuated than under self-guarding. Since immune activation requires both accumulation of the guard signal and subsequent activation of the immune response, rapid fluctuations in the trigger signal are damped at two distinct stages. Second, the additional timescale 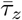 introduces further response delay. When *γ*_off_ ≲ 1, guard turnover occurs on a timescale comparable to regulatory dynamics, and the delay becomes dynamically significant.

It is important to note, however, that the guarded architecture does not necessarily produce a weaker immune response than self-guarding. When *z* ≈ 0, the guard dynamics satisfy equation (15); for trigger signals that vary on timescales slower than 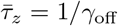, the guard activation then approximately tracks the trigger with gain

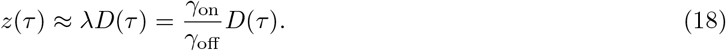

Thus, when the gain factor *λ >* 1, guarding can exhibit enhanced sensitivity to moderate deviations in the trigger signal, with *z* exceeding *D* prior to saturation. Since immune activation is driven by *z* via equation (8), this amplification can translate into a larger (but slower) initial immune response *y* compared with self-guarding under the same trigger signal. The guard activation saturates as *z* → 1 and so amplification primarily affects the early and intermediate phases of activation, when *z* remains small. Guarding therefore introduces two distinct effects: an additional dynamical timescale that increases temporal filtering, and a gain factor *γ*_on_*/γ*_off_ that can enhance sensitivity when *γ*_on_ *> γ*_off_. Whether guarding is overall more conservative than self-guarding therefore depends on the balance between these temporal and gain effects.

Overall, guarding can enhance robustness to perturbations in the regulatory factor by increasing temporal averaging, which can reduce the propensity for false-positive immune responses. However, this robustness comes at the cost of slower activation during genuine infection. Since viral growth is exponential prior to immune control, even modest differences in activation timing may translate into appreciable differences in peak viral load. The net effect of guarding relative to self-guarding therefore depends on parameter balance: while the additional filtering generally increases response latency, guarding can amplify moderate trigger signals when *γ*_on_ *> γ*_off_, potentially leading to a larger, but delayed, immune response.

These findings are illustrated in Figs. 2-4. Self-guarding produces a faster immune response (Fig. 2C, 3C) and lower peak viral load (Fig. 2B, 3B) than guarding provided the gain factor *λ* = *γ*_on_*/γ*_off_ ≤ 1. However, the peak immune response and viral load both depend on the gain factor and the relative rates of the guard activation signal, with slower guard dynamics leading to a higher peak immune response (Fig. 3C) and a gain factor less than one leading to a higher peak viral load (Fig. 2B, 3B).

**Figure 2:**
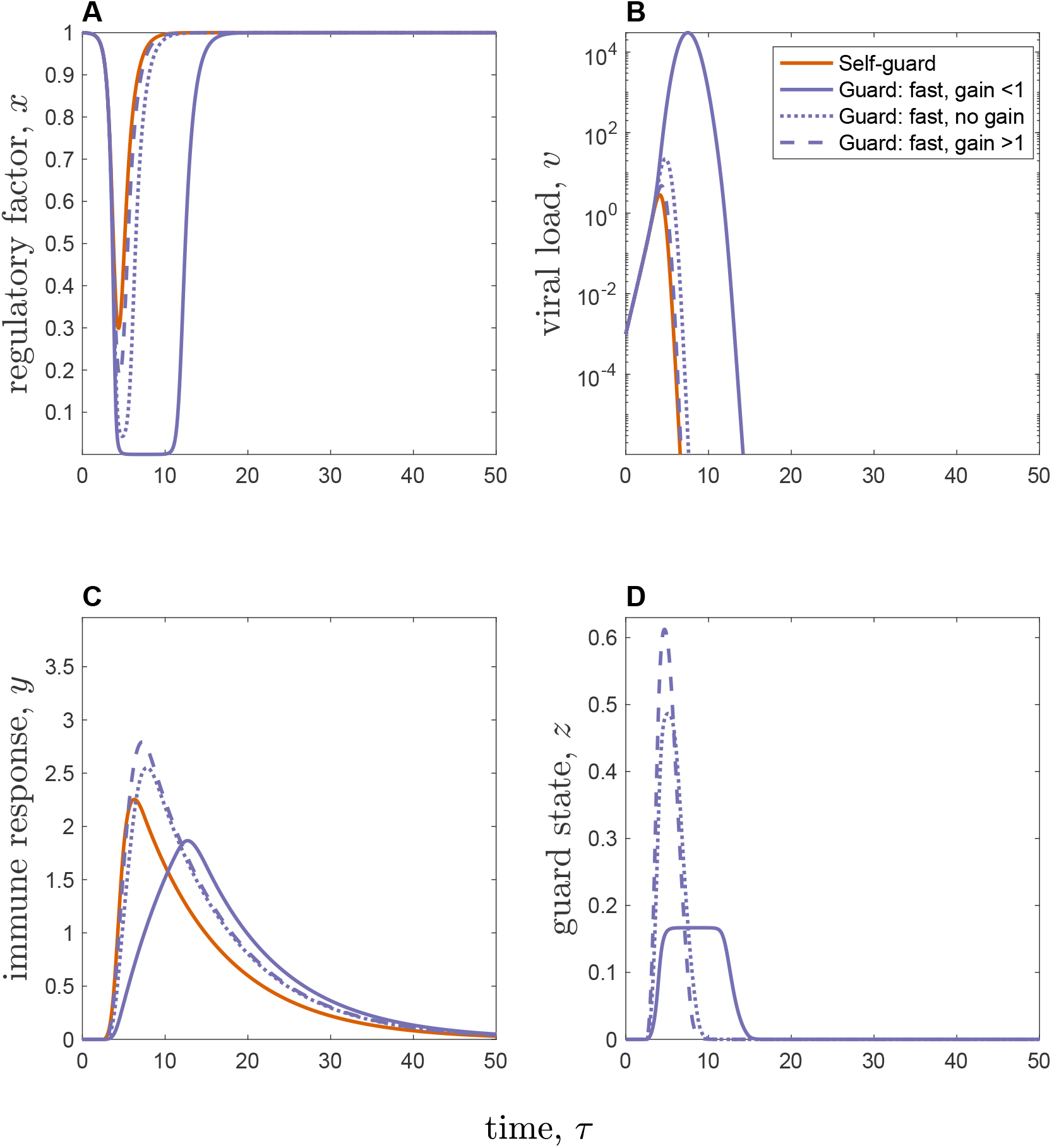
Pathogen and immune dynamics under self-guarding and (fast) guarding. Self-guarding is shown in orange and guarding variants are shown in pr with different line styles corresponding to different gain factors: (i) gain factor *λ <* 1 (*γ*_on_ = 0.5, *γ*_off_ = 2.5; solid); (ii) no gain factor *λ* = 1 (*γ*_on_ = 2.5, *γ*_off_ = 2.5; dotted); (iii) gain factor *λ >* 1 (*γ*_on_ = 5.0, *γ*_off_ = 2.5; dashed). (A) Regulatory factor, *x*. (B) Viral load, *v* (log scale); trajectories are truncated once *v* drops below the clearance threshold *v*_thresh_. (C) Immune response, *y*. (D) Guard state, *z* (guarding only). Other parameters: *ρ* = 5.0, *α* = 0.6, *θ* = 0.95, 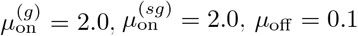.

**Figure 3:**
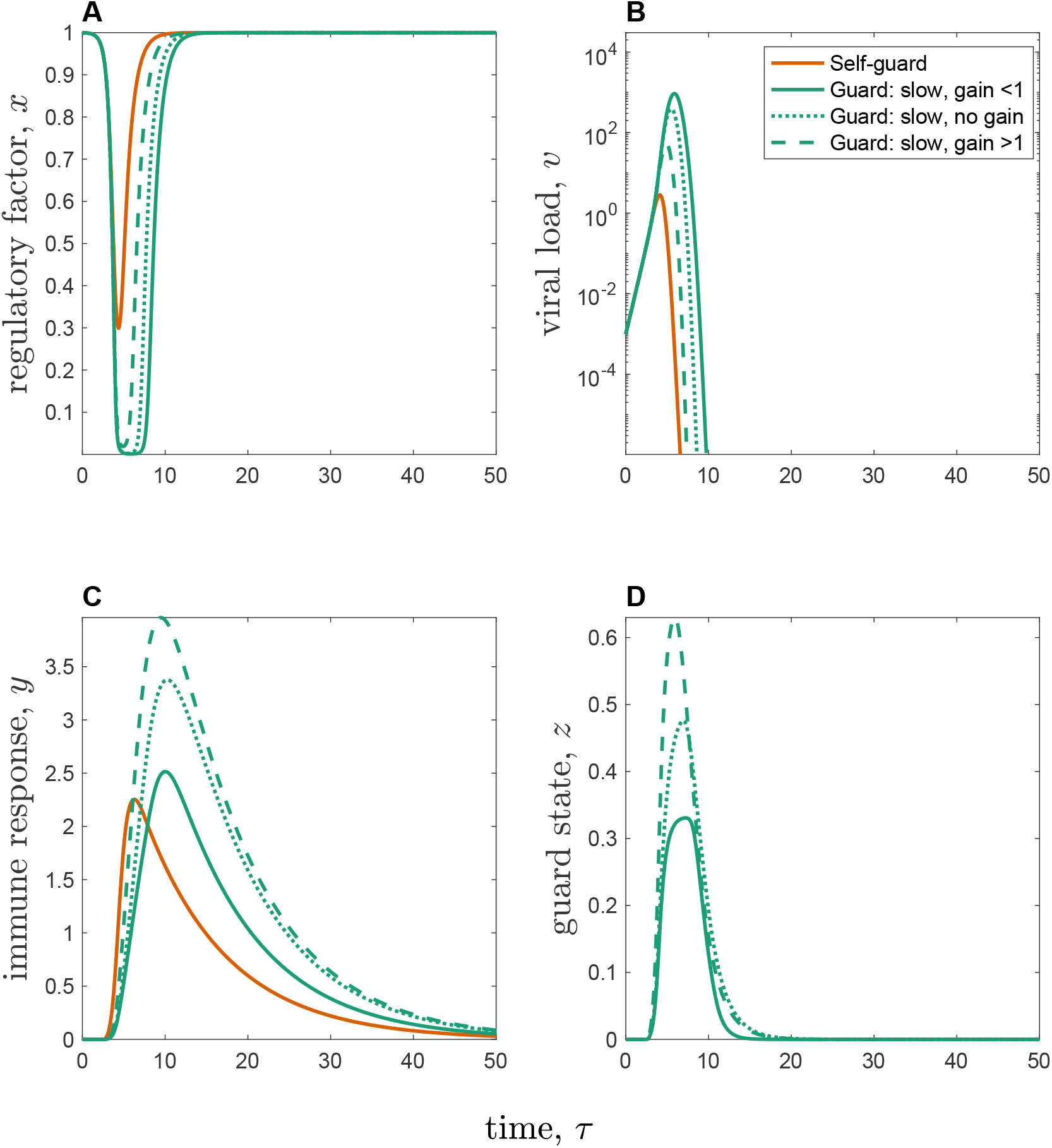
Pathogen and immune dynamics under self-guarding and (slow) guarding. Self-guarding is shown in orange and guarding variants are shown in green with different line styles corresponding to different gain factors: (i) gain factor *λ <* 1 (*γ*_on_ = 0.5, *γ*_off_ = 1.0; solid); (ii) no gain factor *λ* = 1 (*γ*_on_ = 0.5, *γ*_off_ = 0.5; dotted); (iii) gain factor *λ >* 1 (*γ*_on_ = 1.0, *γ*_off_ = 0.5; dashed). (A) Regulatory factor, *x*. (B) Viral load, *v* (log scale); trajectories are truncated once *v* drops below the clearance threshold *v*_thresh_. (C) Immune response, *y*. (D) Guard state, *z* (guarding only). Other parameters as described in Fig. 2.

In the absence of the virus, noise in the regulatory factor can lead to false-positive immune responses (Fig. 4). As self-guarding only operates as a single low-pass filter on the regulatory factor, the potential for false-positive immune responses is high and is less tunable by natural selection (i.e., natural selection would primarily act on the activation threshold, *θ*). However, guarding only notably reduces false-positive immune responses when the gain factor is less than one and this effect is larger when guard dynamics are relatively fast (Fig. 4C-D). Together, these results suggest that guarding is only likely to be favoured over self-guarding if it substantially reduces the false-positive rate. Intuitively, the proportion of time that the regulatory factor falls below the activation threshold increases with noise strength *σ*_*x*_ and decreases with higher triggering thresholds *θ* (Fig. 4B). Note that tuning the triggering threshold alone generally cannot fully compensate for increased stochasticity in the regulatory factor.

**Figure 4:**
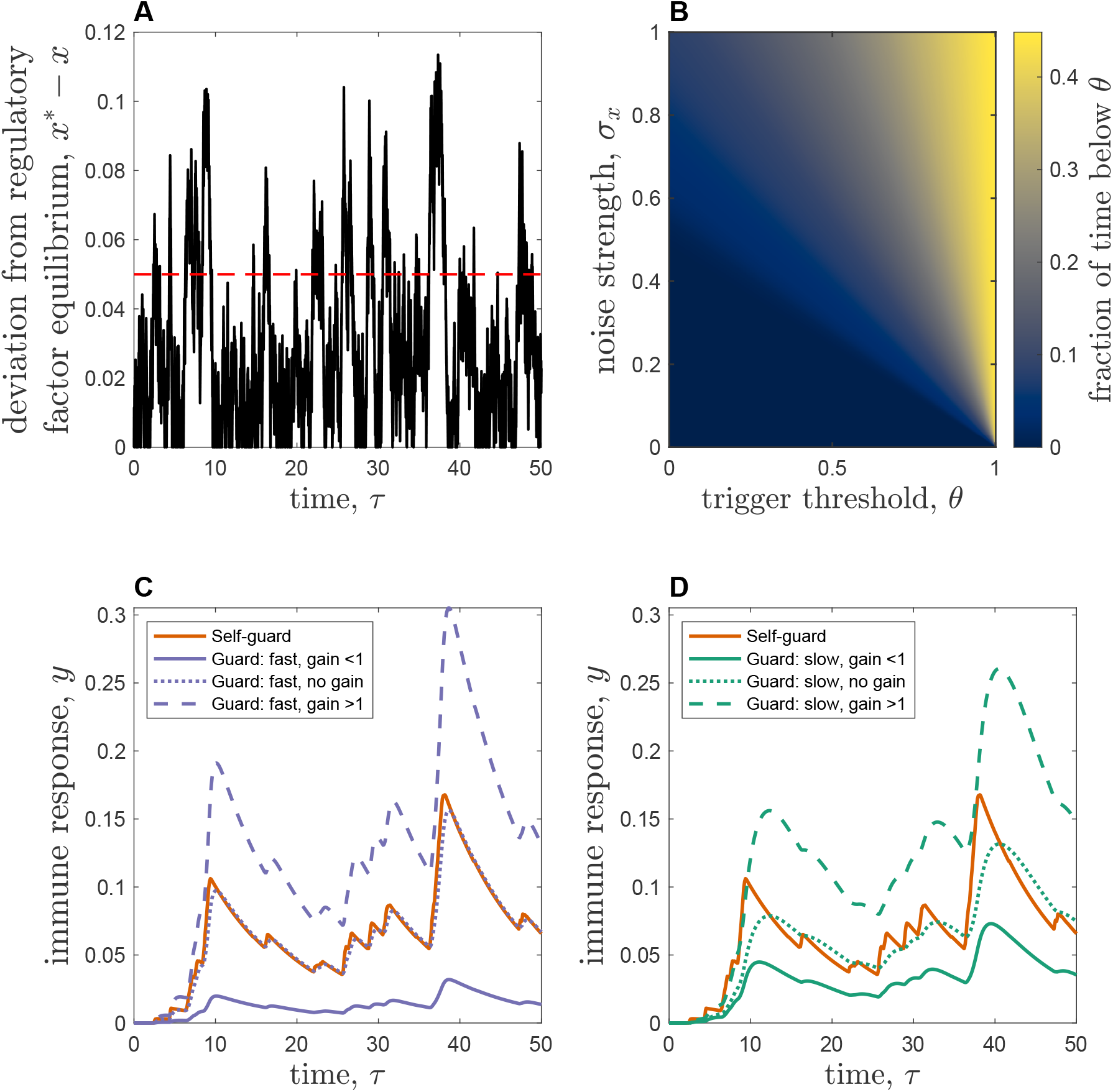
False-positive immune activation under self-guarding and guarding. Stochastic simulations in the absence of the virus (*v* = 0) and additive noise applied only to the regulatory factor *x*. (A) Deviation of the regulatory factor, *x*, from it’s equilibrium, *x*^∗^ = 1, with the activation threshold 1 − *θ* = 0.05 (dashed). The same stochastic trajectory for the regulatory factor signal (*x*) is used across all models. (B) Fraction of time that the regulatory factor *x* lies below the activation threshold *θ* in the absence of infection (*v* = 0), as a function of noise strength *σ*_*x*_ and threshold *θ*. (C-D) Immune response, *y* for self-guarding (orange) with (C) fast (purple) and (D) slow (green) guarding regimes. Gain factors and other parameters as described in Fig. 2-3.

## Discussion

In many host-pathogen systems, natural selection has led to the evolution of multi-layered immune responses, whereby hosts can either directly detect the presence of pathogens or indirectly detect their effects by guarding key pathogen targets to detect modification (Janeway and Medzhitov 2002; Dangl and Jones 2001). Recently, a self-guarding mechanism has been discovered, where a protein carries out both guard and guardee functions as a negative regulator of a secondary immune response (Gaidt et al. 2021). This appears to create a strong constraint on pathogen evolution, as targeting a self-guarding protein may be necessary for replication but will necessarily trigger a secondary immune response. When guard and guardee functions are separated, it is possible for pathogens to also target the guard to evade detection (Cui et al. 2015; Dodds and Rathjen 2010). This raises the question as to when self-guarding through negative regulation should be favoured over regular guarding. Here, we have explored mathematical models of guarding and self-guarding architectures to compare their impact on pathogen and immune dynamics. We have shown that while self-guarding generates a faster immune response that typically leads to lower pathogen load, it is also more at risk of false-positives and therefore may lead to additional auto-immune or metabolic costs.

The key dynamical difference between guarding and self-guarding is that separate guard and guardee functions introduce an additional filtering stage which increases temporal averaging of potentially noisy signals from the guardee. Self-guarding lacks this two-stage filtering and is therefore more responsive, but also more sensitive to noise. However, guarding is only more effective at reducing false-positives in our model if the additional filtering stage has a gain factor of less than one, which means that guard activation responds sublinearly to perturbations in the guarded pathway. Such behaviour is consistent with general features of immune signalling systems. For example, innate immune pathways such as NF-*κ*B signalling are subject to strong negative feedback regulation, including I*κ*B-mediated inhibition, which limits amplification (Hoffmann et al. 2002). Immune signalling pathways also exhibit limited information transmission, such that only a restricted range of input variation can be reliably distinguished at the level of downstream responses (Cheong et al. 2011). An effective gain of less than one for guarded pathways is therefore consistent with other well-established immune signalling pathways.

Although our model is phenomenological, we can infer several qualitative predictions for when self-guarding may be favoured over separate guarding functions. First, changes in the self-guard should be highly specific to pathogen activity rather than more generic processes such as cellular stress, and should produce a large, unambiguous signal. Second, intrinsic fluctuations in the self-guard should be small relative to the immune activation threshold, such that stochastic excursions below the threshold are rare in the absence of infection. Third, the costs of delayed immune activation should be high (e.g., in the case of rapidly replicating or highly virulent pathogens where even modest delays substantially increase peak pathogen load). Under such conditions, the speed advantage of self-guarding may outweigh its increased susceptibility to false-positives. Self-guarding should therefore be favoured when regulatory dynamics are relatively stable and infection produces a strong, specific perturbation. Conversely, self-guarding is unlikely to evolve in regulatory pathways with inherently noisy dynamics, where stochastic fluctuations would frequently cross the activation threshold.

Our model was inspired by the discovery of self-guarding against HSV-1 (Gaidt et al. 2021) and poxvirus (Remick et al. 2026). These systems are broadly consistent with the qualitative predictions of our model. First, in both cases the immune response is triggered by a specific pathogen-driven perturbation rather than a generic consequence of cellular stress, and thus represents an infection-specific trigger. Second, the targeted host factors (e.g., MORC3, N4BP1 and ZC3H12A) act as negative regulators of immune signalling pathways (Gaidt et al. 2021; Remick et al. 2026). Chromatin regulators such as MORC3 typically function within relatively stable nuclear complexes (Hager et al. 2009), and similarly the poxvirus-targeted factors participate in established antiviral regulatory pathways, so that their disruption likely generates a strong signal-to-noise ratio. Third, viruses such as HSV-1 replicate rapidly and can encode multiple immune antagonists (Su et al. 2016; Danastas et al. 2020), which implies that early immune activation could be particularly important for limiting viral growth. These features may help explain why self-guarding has evolved in these systems. Notably, however, the poxvirus example differs in that self-guarding is distributed across multiple host proteins rather than confined to a single protein, demonstrating that self-guarding need not be limited to a single protein. While few examples of self-guarding have been identified compared to guarding, this may simply be due to the fact that this mechanism has only recently been discovered.

We believe our model is the first to compare guarding and self-guarding architectures, but previous theoretical work has shown that other immune sensitivity and activation thresholds can evolve in response to the relative risks of false-positives and infection (McKeithan 1995; Shudo and Iwasa 2001; Frank 2002; Metcalf et al. 2017). More generally, optimal immune activation is predicted to balance the risk of autoimmunity or immunopathology against the risk of uncontrolled infection (Graham et al. 2005; Graham et al. 2022). Evolutionary models show that immune responsiveness is shaped by the costs of erroneous or excessive activation (Shudo and Iwasa 2001). At a mechanistic level, models of T cell receptor signalling show that additional processing steps can enhance discrimination by reducing spurious activation, albeit at the cost of slower response times (McKeithan 1995). More broadly, our results relate to predictions for the evolution of indirect pathogen detection, which suggest that hosts may evolve to sense perturbations of host processes rather than pathogen molecules directly when pathogen evasion of direct recognition is likely (Dangl and Jones 2001; Frank 2002). Furthermore, negative immune regulators are expected to evolve when the costs of excessive or erroneous immune activation are high (Frank and Schmid-Hempel 2019; Frank 2023).

Our results add to this body of work by demonstrating that the architecture of immune guarding can affect the risk of false-positives, and hence the trade-off for more responsive (self-guarding) versus more robust (guarding) secondary immune triggers. Guarding therefore changes the dynamics of immune activation, rather than simply shifting the activation threshold. Furthermore, our results show that while separate guard and guardee functions can reduce false-positives, this only occurs when the gain factor (defined as the quotient of the non-dimensionalised guard activation and deactivation rates, *λ* = *γ*_on_*/γ*_off_) is less than 1.

The model presented herein is a simplified representation of guarding and self-guarding immune pathways as we wished to determine the fundamental differences between these architectures at the initial signal processing stage. Our model therefore only focuses on an “all else being equal” scenario before a secondary immune response is triggered. However, it is possible that self-guarded constituents of host immune pathways could be subject to additional downstream regulation to reduce the risk of false-positives. Self-guarding may also be limited by molecular or immunological constraints of encompassing two distinct functions (e.g., antiviral properties and negative regulation of an immune response) in a single protein. We have neglected other potential costs and benefits of guarding or self-guarding in our model that may shape their evolution. For example, a key additional advantage of self-guarding is that it may be less prone to evolutionary adaptations by pathogens than when guard and guardee functions are separate. For instance, pathogens may evolve to modify or suppress the guard itself, interfere with downstream signalling pathways, or redirect their effectors toward alternative host targets that avoid triggering guarded surveillance (Dodds and Rathjen 2010). However, having a separate guard may also allow for hosts to evolve decoy host proteins, which mimic actual pathogen targets but function primarily as immune sensors (Hoorn and Kamoun 2008; Dodds and Rathjen 2010; Le Roux et al. 2015). Theory suggests that such decoys can be favoured when they improve immune detection while reducing the fitness costs of perturbing functional host targets (Ispolatov and Doebeli 2010).

Understanding how immune architectures shape host and pathogen evolution remains an important goal. Here, we have shown that costs of self-guarding are most likely to arise through a higher risk of false-positive immune activation. Our results therefore suggest that the benefits of self-guarding should be greatest in systems where pathogen-induced perturbations are strong and specific, but background fluctuations in the guarded process are small.

## References

Braud, V. M., D. S. J. Allan, C. A. O’Callaghan, K. Söderström, A. D’Andrea, G. S. Ogg, S. Lazetic, N. T. Young, J. I. Bell, J. H. Phillips, L. L. Lanier, and A. J. McMichael (1998). HLA-E binds to natural killer cell receptors CD94/NKG2A, B and C. Nature 391.6669, pp. 795–799. doi: 10.1038/35869.

Cheong, R., A. Rhee, C. J. Wang, I. Nemenman, and A. Levchenko (2011). Information transduction capacity of noisy biochemical signaling networks. Science 334.6054, pp. 354–358. doi: 10.1126/science.1204553.

Cui, H., K. Tsuda, and J. E. Parker (2015). Effector-triggered immunity: from pathogen perception to robust defense. Annual Review of Plant Biology 66, pp. 487–511. doi: 10.1146/annurev-arplant-050213-040012.

Danastas, K., M. Miranda-Saksena, A. L. Cunningham, and N. K. Saksena (2020). Herpes simplex virus type 1 interactions with the interferon system. International Journal of Molecular Sciences 21.14, p. 5150. doi: 10.3390/ijms21145150.

Dangl, J. L. and J. D. G. Jones (2001). Plant pathogens and integrated defence responses to infection. Nature 411.6839, pp. 826–833. doi: 10.1038/35081161.

Dodds, P. N. and J. P. Rathjen (2010). Plant immunity: towards an integrated view of plant–pathogen interactions. Nature Reviews Genetics 11.8, pp. 539–548. doi: 10.1038/nrg2812.

Frank, S. A. (2002). Immunology and Evolution of Infectious Disease. Princeton, NJ: Princeton University Press.

Frank, S. A. (2023). Disease from opposing forces in regulatory control. Evolution, Medicine, and Public Health 11.1, pp. 348–352. doi: 10.1093/emph/eoad033.

Frank, S. A. and P. Schmid-Hempel (2019). Evolution of negative immune regulators. PLOS Pathogens 15.8, e1007913. doi: 10.1371/journal.ppat.1007913.

Gaidt, M. M., A. Morrow, M. R. Fairgrieve, J. P. Karr, N. Yosef, and R. E. Vance (2021). Self-guarding of MORC3 enables virulence factor-triggered immunity. Nature 600.7887, pp. 138–142. doi: 10.1038/s41586-021-04054-5.

Göhre, V. and S. Robatzek (2008). Breaking the barriers: microbial effector molecules subvert plant immunity. Annual Review of Phytopathology 46, pp. 189–215.

Graham, A. L., J. E. Allen, and A. F. Read (2005). Evolutionary causes and consequences of immunopathology. Annual Review of Ecology, Evolution, and Systematics 36, pp. 373–397. doi: 10.1146/annurev.ecolsys.36.102003.152622.

Graham, A. L., E. C. Schrom, and C. J. E. Metcalf (2022). The evolution of powerful yet perilous immune systems. Trends in Immunology 43.2, pp. 89–101. doi: 10.1016/j.it.2021.11.002.

Hager, G. L., J. G. McNally, and T. Misteli (2009). Transcription dynamics. Molecular Cell 35.6, pp. 741– 753. doi: 10.1016/j.molcel.2009.09.005.

Hansen, T. H. and M. Bouvier (2009). MHC class I antigen presentation: learning from viral evasion strategies. Nature Reviews Immunology 9.7, pp. 503–513. doi: 10.1038/nri2575.

Hoffmann, A., A. Levchenko, M. L. Scott, and D. Baltimore (2002). The IκB-NF-κB signaling module: temporal control and selective gene activation. Science 298.5596, pp. 1241–1245. doi: 10.1126/science.1071914.

Hoorn, R.A.L. van der and S. Kamoun (2008). From Guard to Decoy: A New Model for Perception of Plant Pathogen Effectors. The Plant Cell 20.8, pp. 2009–2017. doi: 10.1105/tpc.108.060194.

Ilchenko, K. and E. Pfeifer (2024). A layered effect in bacterial defense. Cell Host & Microbe 32.4, pp. 447– 449. issn: 1931-3128. doi: 10.1016/j.chom.2024.03.007.

Ispolatov, I. and M. Doebeli (2010). On the Evolution of Decoys in Plant Immune Systems. Biological Theory 5.3, pp. 256–263. doi: 10.1162/BIOT_a_00055.

Janeway, C. A. (1989). Approaching the asymptote? Evolution and revolution in immunology. Cold Spring Harbor Symposia on Quantitative Biology 54, pp. 1–13.

Janeway, C. A. J. and R. Medzhitov (2002). Innate immune recognition. Annual Review of Immunology 20, pp. 197–216. doi: 10.1146/annurev.immunol.20.083001.084359.

Jones, J. D. and J. L. Dangl (2006). The plant immune system. Nature 444, pp. 323–329.

Kloeden, P. E. and E. Platen (1992). Numerical Solution of Stochastic Differential Equations. Vol. 23. Applications of Mathematics. Berlin: Springer. doi: 10.1007/978-3-662-12616-5.

Lamkanfi, M. and V. M. Dixit (2014). Mechanisms and functions of inflammasomes. Cell 157.5, pp. 1013– 1022. doi: 10.1016/j.cell.2014.04.007.

Lanier, L. L. (2005). NK cell recognition. Annual Review of Immunology 23, pp. 225–274. doi: 10.1146/annurev.immunol.23.021704.115526.

Le Roux, C., G. Huet, A. Jauneau, L. Camborde, D. Trémousaygue, A. Kraut, B. Zhou, M. Levaillant, H. Adachi, H. Yoshioka, S. Raffaele, R. Berthomé, Y. Couté, J. E. Parker, and L. Deslandes (2015). A receptor pair with an integrated decoy converts pathogen disabling of transcription factors to immunity. Cell 161.5, pp. 1074–1088. doi: 10.1016/j.cell.2015.04.025.

Lindeberg, M., S. Cunnac, and A. Collmer (2012). Pseudomonas syringae type III effector repertoires: last words in endless arguments. Trends in Microbiology 20.4, pp. 199–208. doi: 10.1016/j.tim.2012.01.003.

Ljunggren, H.-G. and K. Kärre (1990). In search of the ‘missing self’: MHC molecules and NK cell recognition. Immunology Today 11.7, pp. 237–244. doi: 10.1016/0167-5699(90)90097-S.

Mackey, D., B. F. Holt, A. Wiig, and J. L. Dangl (2002). RIN4 interacts with Pseudomonas syringae type III effector molecules and is required for RPM1-mediated resistance in Arabidopsis. Cell 108, pp. 743–754.

McKeithan, T. W. (1995). Kinetic proofreading in T-cell receptor signal transduction. Proceedings of the National Academy of Sciences 92.11, pp. 5042–5046.

Metcalf, C. J. E., A. T. Tate, and A. L. Graham (2017). Demographically framing trade-offs between sensitivity and specificity illuminates selection on immunity. Nature Ecology & Evolution 1, pp. 1766–1772. doi: 10.1038/s41559-017-0315-3.

Netea, M. G., A. Schlitzer, K. Placek, L. A. Joosten, and J. L. Schultze (2019). Innate and Adaptive Immune Memory: an Evolutionary Continuum in the Host’s Response to Pathogens. Cell Host & Microbe 25.1, pp. 13–26. issn: 1931-3128. doi: 10.1016/j.chom.2018.12.006.

Remick, B. C., M. M. Gaidt, and R. E. Vance (2023). Effector-Triggered Immunity. Annual Review of Immunology 41.Volume 41, 2023, pp. 453–481. issn: 1545-3278. doi: 10.1146/annurev-immunol-101721-031732.

Remick, B. C., J. Q. Mao, A. G. Manford, A. D. Gutierrez-Jensen, A. Wagner, M. Rape, G. McFadden, M. M. Rahman, M. M. Gaidt, and R. E. Vance (2026). Poxvirus attack of antiviral defense pathways unleashes an effector-triggered NF-κB response. Science 391.6786, eadw4937. doi: 10.1126/science.adw4937.

Shudo, E. and Y. Iwasa (2001). Inducible defense against pathogens and parasites: optimal choice among multiple options. Journal of Theoretical Biology 209.2, pp. 233–247.

Su, C., G. Zhan, and C. Zheng (2016). Evasion of host antiviral innate immunity by HSV-1, an update. Virology Journal 13, p. 38. doi: 10.1186/s12985-016-0495-5.

Tran, A. X., C. M. Stead, and M. S. Trent (2005). Remodeling of Helicobacter pylori lipopolysaccharide. Journal of Endotoxin Research 11.3, pp. 161–166. doi: 10.1179/096805105X37349.

Van der Biezen, E. A. and J. D. G. Jones (1998). Plant disease-resistance proteins and the gene-for-gene concept. Trends in Biochemical Sciences 23.12, pp. 454–456. doi: 10.1016/S0968-0004(98)01311-5.

Vasu, K. and V. Nagaraja (2013). Diverse functions of restriction-modification systems in bacteria. Nucleic Acids Research 41, pp. 20–35.

Vivier, E., D. H. Raulet, L. Moretta, et al. (2011). Innate or adaptive immunity? The example of natural killer cells. Science 331, pp. 44–49.

Yewdell, J. W. and J. R. Bennink (1999). Mechanisms of viral interference with MHC class I antigen processing and presentation. Annual Review of Cell and Developmental Biology 15, pp. 579–606. doi: 10.1146/annurev.cellbio.15.1.579.

Züst, R., L. Cervantes-Barragan, M. Habjan, R. Maier, B. W. Neuman, J. Ziebuhr, K. J. Szretter, S. C. Baker, W. Barchet, M. S. Diamond, S. G. Siddell, B. Ludewig, and V. Thiel (2011). Ribose 2’-O-methylation provides a molecular signature for the distinction of self and non-self mRNA dependent on the RNA sensor Mda5. Nature Immunology 12.2, pp. 137–143. doi: 10.1038/ni.1979.

