## Supplementary material for "Guarding versus self-guarding in innate immunity": source code: guarding_vs_self_guarding.m

```
%% guarding_vs_self_guarding.m
% Reproduces deterministic and stochastic simulations for the manuscript
% "Guarding versus self-guarding in innate immunity".
%
% Modified to:
%   1. Split deterministic guarded trajectories into two separate figures:
%      one for the fast guardsets and one for the slow guardsets.
%   2. Rearrange the stochastic figure into a 2x2 layout:
%        A: deviation from regulatory-factor equilibrium
%        B: heatmap
%        C: immune response with fast guardsets
%        D: immune response with slow guardsets
%   3. Use colorblindness- and print-safe colors.
%
% Notes:
%   - In stochastic runs, x is clamped to [0,1], y to y >= 0, z to [0,1]
%     for guarded runs, z = 0 for self-guarding, and v is fixed at 0.
%   - This file is self-contained and requires no other .m files.

clear; close all; clc;

%% ------------------- Parameters (non-dimensional) -------------------
p.rho         = 5.0;   % r/s (unused in SDE when v is clamped to 0)
p.alpha       = 0.6;   % dR*R0/r (unused in SDE when v is clamped to 0)
p.theta       = 0.95;  % Ra/R0
p.gamma_on    = 0.5;   % g_on/(s*Gmax)
p.gamma_off   = 2.5;   % g_off/s
p.mu_on_guard = 2.0;   % (dI*m_on*Gmax)/(r*s)
p.mu_on_self  = 2.0;   % (dI*m_on)/(r*s)
p.mu_off      = 0.1;   % m_off/s

% Trigger smoothing parameter for the sqrt-smoothed hinge.
p.c      = 1e-3;

% Additive noise strength for x in false-positive SDE runs.
p.sigma_x     = 0.05;

%% ------------------- Simulation settings -------------------
tauSpan = [0, 50];

% Plot colours (RGB scaled to [0,1])
col_guard_fast = [117, 112, 179] / 255;
col_guard_slow = [27, 158, 119] / 255;
col_self       = [217, 95, 2] / 255;

% Figure position
figPos = [500   250   700   700];

% Guard parameter sets to compare
% Ordered so legends read:
%   gain < 1, no gain, gain > 1
guardSets = [ ...
    struct('gamma_off',2.5,'gamma_on',0.5,'ls','-','color',col_guard_fast,'speed','fast','label','Guard: fast, gain <1'); ...
    struct('gamma_off',2.5,'gamma_on',2.5,'ls',':','color',col_guard_fast,'speed','fast','label','Guard: fast, no gain'); ...
    struct('gamma_off',2.5,'gamma_on',5.0,'ls','--','color',col_guard_fast,'speed','fast','label','Guard: fast, gain >1'); ...
    struct('gamma_off',1.0,'gamma_on',0.5,'ls','-','color',col_guard_slow,'speed','slow','label','Guard: slow, gain <1'); ...
    struct('gamma_off',0.5,'gamma_on',0.5,'ls',':','color',col_guard_slow,'speed','slow','label','Guard: slow, no gain'); ...
    struct('gamma_off',0.5,'gamma_on',1.0,'ls','--','color',col_guard_slow,'speed','slow','label','Guard: slow, gain >1') ...
];

% Viral clearance threshold for deterministic runs.
% Once v crosses below v_thresh, we treat the virus as cleared and enforce
% an absorbing state v \equiv 0 for the remainder of the simulation.
v_thresh = 1e-6;

% Initial conditions
% State vector is Y = [x; v; y; z].
y0_ode = [1; 1e-3; 0; 0];

y0_sde = y0_ode;
y0_sde(2) = 0;  % false-positive runs start with v = 0

% ODE solver tolerances
odeOpts = odeset('RelTol',1e-8,'AbsTol',1e-10);

% SDE step and RNG (Euler-Maruyama)
dt = 1e-3;
rng(1);  % reproducible shared noise trajectory

% Clamp stochastic trajectories to biologically meaningful bounds.
doClamp = true;

%% ===================== 1) Deterministic ODE =========================
% Deterministic infection trajectories with an absorbing cleared-virus rule
% to prevent rebounds once v falls below v_thresh.

% Self-guarding
[tS_ode, YS_ode] = solve_ode_absorbing_clearance( ...
    @guard_models_nd_sqrt, tauSpan, y0_ode, odeOpts, p, "self", v_thresh);

% Guarding for six parameter sets
ntypes = numel(guardSets);
tG_ode = cell(ntypes,1);
YG_ode = cell(ntypes,1);
for k = 1:ntypes
    pk = p;
    pk.gamma_off = guardSets(k).gamma_off;
    pk.gamma_on  = guardSets(k).gamma_on;
    [tG_ode{k}, YG_ode{k}] = solve_ode_absorbing_clearance( ...
        @guard_models_nd_sqrt, tauSpan, y0_ode, odeOpts, pk, "guarded", v_thresh);
end

% Global axis limits so the two deterministic figures are directly comparable
odeLims = compute_ode_global_limits(tS_ode, YS_ode, tG_ode, YG_ode, v_thresh);

% Fast guardsets
plot_figure2_ode_group( ...
    tS_ode, YS_ode, tG_ode(1:3), YG_ode(1:3), guardSets(1:3), ...
    v_thresh, col_self, odeLims, figPos);

% Slow guardsets
plot_figure2_ode_group( ...
    tS_ode, YS_ode, tG_ode(4:6), YG_ode(4:6), guardSets(4:6), ...
    v_thresh, col_self, odeLims, figPos);

%% ===================== 2) Stochastic SDE (false positives) ==========
% Pre-generate a common x-noise trajectory (Wiener increments) and reuse it
% in both architectures so that differences are structural.
tau0 = tauSpan(1);
tauF = tauSpan(2);
N = floor((tauF - tau0)/dt) + 1;
dW_x = sqrt(dt) * randn(N-1, 1);

% Self-guarding
[tS_sde, YS_sde] = simulate_em_common_xnoise( ...
    @guard_models_nd_sde_falsepos_xnoise, tauSpan, y0_sde, dt, p, "self", doClamp, dW_x);

% Guarding for six parameter sets
tG_sde = cell(ntypes,1);
YG_sde = cell(ntypes,1);
for k = 1:ntypes
    pk = p;
    pk.gamma_off = guardSets(k).gamma_off;
    pk.gamma_on  = guardSets(k).gamma_on;
    [tG_sde{k}, YG_sde{k}] = simulate_em_common_xnoise( ...
        @guard_models_nd_sde_falsepos_xnoise, tauSpan, y0_sde, dt, pk, "guarded", doClamp, dW_x);
end

% Heatmap settings for the x-only false-positive process:
%   dx = (1 - x) dt + sigma_x dW
heatmapCfg.sigma_vals  = linspace(0, 1, 500);
heatmapCfg.theta_vals  = linspace(0, 1, 500);
heatmapCfg.burninTime  = 10;

frac_below = compute_frac_below_heatmap( ...
    tauSpan, dt, dW_x, heatmapCfg.sigma_vals, heatmapCfg.theta_vals, heatmapCfg.burninTime);

plot_figure3_sde_2x2( ...
    tS_sde, YS_sde, ...
    tG_sde(1:3), YG_sde(1:3), guardSets(1:3), ...
    tG_sde(4:6), YG_sde(4:6), guardSets(4:6), ...
    col_self, p.theta, ...
    heatmapCfg.theta_vals, heatmapCfg.sigma_vals, frac_below, figPos);

plot_hinge_activation_figure(p.c, p.theta);

%% ====================================================================
%% Local functions used by the deterministic and stochastic simulations
%% ====================================================================

function dydtau = guard_models_nd_sqrt(~, Y, p, modelFlag)
% Right-hand side for the deterministic non-dimensional model with a
% sqrt-smoothed trigger:
%   u = 1 - x/theta
%   D = 0.5*(u + sqrt(u^2 + c^2))

flag = parseFlag(modelFlag);

x = Y(1);
v = Y(2);
y = Y(3);
z = Y(4);

u = 1 - x / p.theta;
D = 0.5 * (u + sqrt(u*u + p.c*p.c));

dx = (1 - x) - x * v;
dv = p.rho * v * (1 - p.alpha * x - y);

switch flag
    case 1
        dz = p.gamma_on * D * (1 - z) - p.gamma_off * z;
        dy = p.mu_on_guard * z - p.mu_off * y;
    case 2
        dz = 0;
        dy = p.mu_on_self * D - p.mu_off * y;
    otherwise
        error('modelFlag must be 1/''guarded'' or 2/''self''.');
end

dydtau = [dx; dv; dy; dz];
end

function [tOut, YOut] = solve_ode_absorbing_clearance(rhsFun, tauSpan, y0, odeOpts, p, modelFlag, v_thresh)
% Solve the deterministic model until v falls below v_thresh, then continue
% with v held at 0 as an absorbing cleared-virus state.

odeOpts1 = odeset(odeOpts, 'NonNegative', 1:4, ...
    'Events', @(t,Y) evt_v_clear(t, Y, v_thresh));
[t1, Y1, te, Ye] = ode45(@(t,Y) rhsFun(t, Y, p, modelFlag), tauSpan, y0, odeOpts1);

if isempty(te)
    tOut = t1;
    YOut = Y1;
    return;
end

t_clear = te(end);
Y_clear = Ye(end, :)';
Y_clear(2) = 0;

if t_clear >= tauSpan(2)
    tOut = t1;
    YOut = Y1;
    YOut(end,2) = 0;
    return;
end

odeOpts2 = odeset(odeOpts, 'NonNegative', 1:4);
[t2, Y2] = ode45(@(t,Y) rhs_cleared(t, Y, p, modelFlag), [t_clear, tauSpan(2)], Y_clear, odeOpts2);

if numel(t2) > 1
    t2 = t2(2:end);
    Y2 = Y2(2:end,:);
end

tOut = [t1; t2];
YOut = [Y1; Y2];

idx_after = find(tOut >= t_clear);
YOut(idx_after,2) = 0;
end

function dY = rhs_cleared(~, Y, p, modelFlag)
% Deterministic dynamics after viral clearance, with v fixed at 0.

Yc = Y;
Yc(2) = 0;
dY = guard_models_nd_sqrt([], Yc, p, modelFlag);
dY(2) = 0;
end

function [value, isterminal, direction] = evt_v_clear(~, Y, v_thresh)
% Event function for first downward crossing of the viral clearance threshold.

v = Y(2);
value = v - v_thresh;
isterminal = 1;
direction = -1;
end

function [f, G] = guard_models_nd_sde_falsepos_xnoise(~, Y, p, modelFlag)
% Drift and diffusion for the false-positive SDE experiment.
% Here v is fixed at 0 and only x has additive noise.

flag = parseFlag(modelFlag);

x = Y(1);
y = Y(3);
z = Y(4);

u = 1 - x / p.theta;
D = 0.5 * (u + sqrt(u*u + p.c*p.c));

dx = (1 - x);
dv = 0;

switch flag
    case 1
        dz = p.gamma_on * D * (1 - z) - p.gamma_off * z;
        dy = p.mu_on_guard * z - p.mu_off * y;
    case 2
        dz = 0;
        dy = p.mu_on_self * D - p.mu_off * y;
    otherwise
        error('modelFlag must be 1/''guarded'' or 2/''self''.');
end

f = [dx; dv; dy; dz];
G = diag([p.sigma_x, 0, 0, 0]);
end

function [tau, Y] = simulate_em_common_xnoise(sdeFun, tauSpan, y0, dt, p, modelFlag, doClamp, dW_x)
% Euler-Maruyama simulation using a supplied x-noise trajectory.

tau0 = tauSpan(1);
tauF = tauSpan(2);
N = floor((tauF - tau0)/dt) + 1;

if numel(dW_x) ~= (N-1)
    error('dW_x must have length N-1 (%d). Got %d.', N-1, numel(dW_x));
end

tau = tau0 + (0:N-1)' * dt;
Y = zeros(N, numel(y0));
Y(1,:) = y0(:)';

flag = parseFlag(modelFlag);

[~, G0] = sdeFun(tau(1), y0(:), p, modelFlag);
m = size(G0, 2);

for n = 1:N-1
    yn = Y(n,:)';
    tn = tau(n);

    [f, G] = sdeFun(tn, yn, p, modelFlag);

    dW = zeros(m,1);
    dW(1) = dW_x(n);
    yn1 = yn + f*dt + G*dW;

    yn1(2) = 0;  % enforce v \equiv 0

    if doClamp
        yn1(1) = min(max(yn1(1), 0), 1);  % x in [0,1]
        yn1(3) = max(yn1(3), 0);          % y >= 0

        if flag == 1
            yn1(4) = min(max(yn1(4), 0), 1);  % z in [0,1]
        else
            yn1(4) = 0;                       % z = 0 for self-guarding
        end
    end

    Y(n+1,:) = yn1';
end
end

function flag = parseFlag(modelFlag)
% Convert model labels to an internal numeric flag.
%   1 = guarded
%   2 = self-guarding

if ischar(modelFlag) || isstring(modelFlag)
    mf = lower(string(modelFlag));

    if mf == "guarded"
        flag = 1;
    elseif mf == "self" || mf == "self-guarding" || mf == "selfguarding"
        flag = 2;
    else
        error('Unknown modelFlag: %s. Use ''guarded'' or ''self''.', string(modelFlag));
    end

elseif isnumeric(modelFlag) && isscalar(modelFlag) && any(modelFlag == [1, 2])
    flag = modelFlag;

else
    error('modelFlag must be 1, 2, ''guarded'', or ''self''.');
end
end

function odeLims = compute_ode_global_limits(tS, YS, tG_cell, YG_cell, v_thresh)
% Compute global axis limits across all deterministic trajectories so the
% fast and slow figures use identical scales.

allT = tS(:);
x_all = YS(:,1);
y_all = YS(:,3);
z_all = [];
v_all = truncate_v_local(YS(:,2), v_thresh);

for k = 1:numel(tG_cell)
    allT = [allT; tG_cell{k}(:)]; %#ok<AGROW>
    x_all = [x_all; YG_cell{k}(:,1)]; %#ok<AGROW>
    y_all = [y_all; YG_cell{k}(:,3)]; %#ok<AGROW>
    z_all = [z_all; YG_cell{k}(:,4)]; %#ok<AGROW>
    v_all = [v_all; truncate_v_local(YG_cell{k}(:,2), v_thresh)]; %#ok<AGROW>
end

v_all = v_all(~isnan(v_all) & v_all > 0);

odeLims.x  = [min(allT), max(allT)];
odeLims.x1 = [min(x_all), max(x_all)];
odeLims.y  = [min(y_all), max(y_all)];
odeLims.z  = [min(z_all), max(z_all)];

if isempty(v_all)
    odeLims.v = [v_thresh, 1];
else
    odeLims.v = [min(v_all), max(v_all)];
end
end

function vplot = truncate_v_local(v, v_thresh)
% Helper for viral-load plotting.
vplot = v;
idx = find(v < v_thresh, 1, 'first');
if ~isempty(idx)
    vplot(idx:end) = NaN;
end
end

function plot_figure2_ode_group(tS, YS, tG_cell, YG_cell, guardSets, v_thresh, col_self, odeLims, figPos)
% Plot deterministic time series for self-guarding and one speed group of
% guarded parameter sets (either fast or slow), using shared axis limits.

figure('Color','w', 'Position', figPos);
ax = gobjects(4,1);

    function vplot = truncate_v(v)
        vplot = v;
        idx = find(v < v_thresh, 1, 'first');
        if ~isempty(idx)
            vplot(idx:end) = NaN;
        end
    end

ax(1) = subplot(2,2,1); hold on;
plot(tS, YS(:,1), '-', 'LineWidth', 2, 'Color', col_self);
for k = 1:numel(guardSets)
    plot(tG_cell{k}, YG_cell{k}(:,1), ...
        'LineStyle', guardSets(k).ls, 'LineWidth', 2, ...
        'Color', guardSets(k).color);
end
set(gca,'FontSize',12,'Box','on');
ylabel('regulatory factor, $x$','Interpreter','latex','FontSize',20);

ax(2) = subplot(2,2,2); hold on;
hSv = plot(tS, truncate_v(YS(:,2)), '-', 'LineWidth', 2, 'Color', col_self);
hGv = gobjects(numel(guardSets),1);
for k = 1:numel(guardSets)
    hGv(k) = plot(tG_cell{k}, truncate_v(YG_cell{k}(:,2)), ...
        'LineStyle', guardSets(k).ls, 'LineWidth', 2, ...
        'Color', guardSets(k).color);
end
set(gca,'YScale','log','FontSize',12,'Box','on');
ylim([0.99*1e-5,1e5])
ylabel('viral load, $v$','Interpreter','latex','FontSize',20);

ax(3) = subplot(2,2,3); hold on;
plot(tS, YS(:,3), '-', 'LineWidth', 2, 'Color', col_self);
for k = 1:numel(guardSets)
    plot(tG_cell{k}, YG_cell{k}(:,3), ...
        'LineStyle', guardSets(k).ls, 'LineWidth', 2, ...
        'Color', guardSets(k).color);
end
set(gca,'FontSize',12,'Box','on');
ylabel('immune response, $y$','Interpreter','latex','FontSize',20);

ax(4) = subplot(2,2,4); hold on;
for k = 1:numel(guardSets)
    plot(tG_cell{k}, YG_cell{k}(:,4), ...
        'LineStyle', guardSets(k).ls, 'LineWidth', 2, ...
        'Color', guardSets(k).color);
end
set(gca,'FontSize',12,'Box','on');
ylabel('guard state, $z$','Interpreter','latex','FontSize',20);

legend(ax(2), [hSv, hGv(:)'], ...
    [{'Self-guard'}, {guardSets.label}], ...
    'FontSize',12,'Location','best','Box','on');

set(ax, 'XLim', odeLims.x);
set(ax(1), 'YLim', odeLims.x1);
set(ax(2), 'YLim', odeLims.v);
set(ax(3), 'YLim', odeLims.y);
set(ax(4), 'YLim', odeLims.z);

linkaxes(ax,'x');

labels = {'A','B','C','D'};
for i = 1:4
    axes(ax(i));
    text(0, 1.01, labels{i}, 'Units','normalized', ...
        'FontWeight','bold', 'FontSize',16, ...
        'HorizontalAlignment','left', 'VerticalAlignment','bottom');
end

annotation(gcf,'textbox',[0 0.01 1 0.05], ...
    'String','time, $\tau$', ...
    'Interpreter','latex', ...
    'HorizontalAlignment','center', ...
    'VerticalAlignment','middle', ...
    'EdgeColor','none', ...
    'FontSize',20);
end

function plot_figure3_sde_2x2(tS, YS, ...
    tG_fast, YG_fast, guardFast, ...
    tG_slow, YG_slow, guardSlow, ...
    col_self, theta, theta_vals, sigma_vals, frac_below, figPos)
% Plot stochastic false-positive trajectories in a 2x2 layout:
%   A: deviation from regulatory-factor equilibrium
%   B: heatmap
%   C: immune response with fast guardsets
%   D: immune response with slow guardsets

figure('Color','w', 'Position', figPos);
ax = gobjects(4,1);

% Shared x-limits for time-series panels
allT = tS(:);
for k = 1:numel(tG_fast)
    allT = [allT; tG_fast{k}(:)]; %#ok<AGROW>
end
for k = 1:numel(tG_slow)
    allT = [allT; tG_slow{k}(:)]; %#ok<AGROW>
end
xlims = [min(allT), max(allT)];

% Shared y-limits for the two immune-response panels
y_all = YS(:,3);
for k = 1:numel(tG_fast)
    y_all = [y_all; YG_fast{k}(:,3)]; %#ok<AGROW>
end
for k = 1:numel(tG_slow)
    y_all = [y_all; YG_slow{k}(:,3)]; %#ok<AGROW>
end
ylims_y = [min(y_all), max(y_all)];

% ---------------- Panel A: x* - x ----------------
ax(1) = subplot(2,2,1); hold(ax(1), 'on');
plot(tS, 1 - YS(:,1), '-', 'LineWidth', 1.5, 'Color', 'k');
plot([0, tS(end)], 1 - theta*[1, 1], '--', 'LineWidth', 1.5, 'Color', 'r');
set(gca,'FontSize',12,'Box','on');
xlabel('time, $\tau$','Interpreter','latex','FontSize',18);
ylabel({'deviation from regulatory','factor equilibrium, $x^{*}-x$'}, ...
    'Interpreter','latex','FontSize',20);

% ---------------- Panel B: heatmap ----------------
ax(2) = subplot(2,2,2);
imagesc(theta_vals, sigma_vals, frac_below);
set(gca, 'YDir', 'normal', 'FontSize', 12, 'LineWidth', 1, 'Box', 'on');
xlabel('trigger threshold, $\theta$','Interpreter','latex','FontSize',18);
ylabel('noise strength, $\sigma_x$','Interpreter','latex','FontSize',20);
ylim([min(sigma_vals), max(sigma_vals)]);
colormap(ax(2), guarding_heatmap_colormap());
cb = colorbar(ax(2));
cb.FontSize = 12;
cb.Ticks = 0:0.1:1;
ylabel(cb, 'fraction of time below $\theta$', ...
    'Interpreter', 'latex', 'FontSize', 18);

% ---------------- Panel C: immune response, fast ----------------
ax(3) = subplot(2,2,3); hold(ax(3), 'on');
hS_fast = plot(tS, YS(:,3), '-', 'LineWidth', 2, 'Color', col_self);
hG_fast = gobjects(numel(guardFast),1);
for k = 1:numel(guardFast)
    hG_fast(k) = plot(tG_fast{k}, YG_fast{k}(:,3), ...
        'LineStyle', guardFast(k).ls, ...
        'LineWidth', 2, ...
        'Color', guardFast(k).color);
end
set(gca,'FontSize',12,'Box','on');
xlabel('time, $\tau$','Interpreter','latex','FontSize',18);
ylabel('immune response, $y$','Interpreter','latex','FontSize',20);
legend([hS_fast, hG_fast(:)'], ...
    [{'Self-guard'}, {guardFast.label}], ...
    'FontSize',10,'Location','northwest','Box','on');

% ---------------- Panel D: immune response, slow ----------------
ax(4) = subplot(2,2,4); hold(ax(4), 'on');
hS_slow = plot(tS, YS(:,3), '-', 'LineWidth', 2, 'Color', col_self);
hG_slow = gobjects(numel(guardSlow),1);
for k = 1:numel(guardSlow)
    hG_slow(k) = plot(tG_slow{k}, YG_slow{k}(:,3), ...
        'LineStyle', guardSlow(k).ls, ...
        'LineWidth', 2, ...
        'Color', guardSlow(k).color);
end
set(gca,'FontSize',12,'Box','on');
xlabel('time, $\tau$','Interpreter','latex','FontSize',18);
ylabel('immune response, $y$','Interpreter','latex','FontSize',20);
legend([hS_slow, hG_slow(:)'], ...
    [{'Self-guard'}, {guardSlow.label}], ...
    'FontSize',10,'Location','northwest','Box','on');

% Shared limits
set(ax([1,3,4]), 'XLim', xlims);
set(ax(3), 'YLim', ylims_y);
set(ax(4), 'YLim', ylims_y);
linkaxes(ax([1,3,4]), 'x');

% Panel labels
labels = {'A','B','C','D'};
for i = 1:4
    axes(ax(i));
    text(0, 1.01, labels{i}, 'Units','normalized', ...
        'FontWeight','bold', 'FontSize',16, ...
        'HorizontalAlignment','left', 'VerticalAlignment','bottom');
end
end

function frac_below = compute_frac_below_heatmap(tauSpan, dt, dW_x, sigma_vals, theta_vals, burninTime)
% Compute frac_below(i,j) = fraction of post-burn-in time for which the
% x-only process
%   dx = (1 - x) dt + sigma_x dW
% lies below theta_vals(j), using one trajectory per sigma_x and then
% evaluating all theta values on that trajectory without resimulation.

tau0 = tauSpan(1);
tauF = tauSpan(2);
N = floor((tauF - tau0)/dt) + 1;

if numel(dW_x) ~= (N-1)
    error('dW_x must have length N-1 (%d). Got %d.', N-1, numel(dW_x));
end

burninIdx = floor(burninTime/dt) + 1;
burninIdx = min(max(burninIdx,1),N);

nSigma = numel(sigma_vals);
nTheta = numel(theta_vals);
frac_below = zeros(nSigma, nTheta);

for i = 1:nSigma
    sigma_x = sigma_vals(i);

    % Simulate one x trajectory for this sigma_x
    x = zeros(N,1);
    x(1) = 1;  % start at homeostasis

    for n = 1:N-1
        x(n+1) = x(n) + (1 - x(n))*dt + sigma_x*dW_x(n);
    end

    % Remove burn-in and evaluate all thresholds at once
    x_ss = x(burninIdx:end);
    below_mat = x_ss < theta_vals;   % implicit expansion
    frac_below(i,:) = mean(below_mat, 1);
end
end

function cm = guarding_heatmap_colormap()
% Custom colormap for the false-positive occupancy heatmap.

cm = [ ...
0.0000 0.1262 0.3015
0.0000 0.1292 0.3077
0.0000 0.1321 0.3142
0.0000 0.1350 0.3205
0.0000 0.1379 0.3269
0.0000 0.1408 0.3334
0.0000 0.1437 0.3400
0.0000 0.1465 0.3467
0.0000 0.1492 0.3537
0.0000 0.1519 0.3606
0.0000 0.1546 0.3676
0.0000 0.1574 0.3746
0.0000 0.1601 0.3817
0.0000 0.1629 0.3888
0.0000 0.1657 0.3960
0.0000 0.1685 0.4031
0.0000 0.1714 0.4102
0.0000 0.1743 0.4172
0.0000 0.1773 0.4241
0.0000 0.1798 0.4307
0.0000 0.1817 0.4347
0.0000 0.1834 0.4363
0.0000 0.1852 0.4368
0.0000 0.1872 0.4368
0.0000 0.1901 0.4365
0.0000 0.1930 0.4361
0.0000 0.1958 0.4356
0.0000 0.1987 0.4349
0.0000 0.2015 0.4343
0.0000 0.2044 0.4336
0.0000 0.2073 0.4329
0.0055 0.2101 0.4322
0.0236 0.2130 0.4314
0.0416 0.2158 0.4308
0.0576 0.2187 0.4301
0.0710 0.2215 0.4293
0.0827 0.2244 0.4287
0.0932 0.2272 0.4280
0.1030 0.2300 0.4274
0.1120 0.2329 0.4268
0.1204 0.2357 0.4262
0.1283 0.2385 0.4256
0.1359 0.2414 0.4251
0.1431 0.2442 0.4245
0.1500 0.2470 0.4241
0.1566 0.2498 0.4236
0.1630 0.2526 0.4232
0.1692 0.2555 0.4228
0.1752 0.2583 0.4224
0.1811 0.2611 0.4220
0.1868 0.2639 0.4217
0.1923 0.2667 0.4214
0.1977 0.2695 0.4212
0.2030 0.2723 0.4209
0.2082 0.2751 0.4207
0.2133 0.2780 0.4205
0.2183 0.2808 0.4204
0.2232 0.2836 0.4203
0.2281 0.2864 0.4202
0.2328 0.2892 0.4201
0.2375 0.2920 0.4200
0.2421 0.2948 0.4200
0.2466 0.2976 0.4200
0.2511 0.3004 0.4201
0.2556 0.3032 0.4201
0.2599 0.3060 0.4202
0.2643 0.3088 0.4203
0.2686 0.3116 0.4205
0.2728 0.3144 0.4206
0.2770 0.3172 0.4208
0.2811 0.3200 0.4210
0.2853 0.3228 0.4212
0.2894 0.3256 0.4215
0.2934 0.3284 0.4218
0.2974 0.3312 0.4221
0.3014 0.3340 0.4224
0.3054 0.3368 0.4227
0.3093 0.3396 0.4231
0.3132 0.3424 0.4236
0.3170 0.3453 0.4240
0.3209 0.3481 0.4244
0.3247 0.3509 0.4249
0.3285 0.3537 0.4254
0.3323 0.3565 0.4259
0.3361 0.3593 0.4264
0.3398 0.3622 0.4270
0.3435 0.3650 0.4276
0.3472 0.3678 0.4282
0.3509 0.3706 0.4288
0.3546 0.3734 0.4294
0.3582 0.3763 0.4302
0.3619 0.3791 0.4308
0.3655 0.3819 0.4316
0.3691 0.3848 0.4322
0.3727 0.3876 0.4331
0.3763 0.3904 0.4338
0.3798 0.3933 0.4346
0.3834 0.3961 0.4355
0.3869 0.3990 0.4364
0.3905 0.4018 0.4372
0.3940 0.4047 0.4381
0.3975 0.4075 0.4390
0.4010 0.4104 0.4400
0.4045 0.4132 0.4409
0.4080 0.4161 0.4419
0.4114 0.4189 0.4430
0.4149 0.4218 0.4440
0.4183 0.4247 0.4450
0.4218 0.4275 0.4462
0.4252 0.4304 0.4473
0.4286 0.4333 0.4485
0.4320 0.4362 0.4496
0.4354 0.4390 0.4508
0.4388 0.4419 0.4521
0.4422 0.4448 0.4534
0.4456 0.4477 0.4547
0.4489 0.4506 0.4561
0.4523 0.4535 0.4575
0.4556 0.4564 0.4589
0.4589 0.4593 0.4604
0.4622 0.4622 0.4620
0.4656 0.4651 0.4635
0.4689 0.4680 0.4650
0.4722 0.4709 0.4665
0.4756 0.4738 0.4679
0.4790 0.4767 0.4691
0.4825 0.4797 0.4701
0.4861 0.4826 0.4707
0.4897 0.4856 0.4714
0.4934 0.4886 0.4719
0.4971 0.4915 0.4723
0.5008 0.4945 0.4727
0.5045 0.4975 0.4730
0.5083 0.5005 0.4732
0.5121 0.5035 0.4734
0.5158 0.5065 0.4736
0.5196 0.5095 0.4737
0.5234 0.5125 0.4738
0.5272 0.5155 0.4739
0.5310 0.5186 0.4739
0.5349 0.5216 0.4738
0.5387 0.5246 0.4739
0.5425 0.5277 0.4738
0.5464 0.5307 0.4736
0.5502 0.5338 0.4735
0.5541 0.5368 0.4733
0.5579 0.5399 0.4732
0.5618 0.5430 0.4729
0.5657 0.5461 0.4727
0.5696 0.5491 0.4723
0.5735 0.5522 0.4720
0.5774 0.5553 0.4717
0.5813 0.5584 0.4714
0.5852 0.5615 0.4709
0.5892 0.5646 0.4705
0.5931 0.5678 0.4701
0.5970 0.5709 0.4696
0.6010 0.5740 0.4691
0.6050 0.5772 0.4685
0.6089 0.5803 0.4680
0.6129 0.5835 0.4673
0.6168 0.5866 0.4668
0.6208 0.5898 0.4662
0.6248 0.5929 0.4655
0.6288 0.5961 0.4649
0.6328 0.5993 0.4641
0.6368 0.6025 0.4632
0.6408 0.6057 0.4625
0.6449 0.6089 0.4617
0.6489 0.6121 0.4609
0.6529 0.6153 0.4600
0.6570 0.6185 0.4591
0.6610 0.6217 0.4583
0.6651 0.6250 0.4573
0.6691 0.6282 0.4562
0.6732 0.6315 0.4553
0.6773 0.6347 0.4543
0.6813 0.6380 0.4532
0.6854 0.6412 0.4521
0.6895 0.6445 0.4511
0.6936 0.6478 0.4499
0.6977 0.6511 0.4487
0.7018 0.6544 0.4475
0.7060 0.6577 0.4463
0.7101 0.6610 0.4450
0.7142 0.6643 0.4437
0.7184 0.6676 0.4424
0.7225 0.6710 0.4409
0.7267 0.6743 0.4396
0.7308 0.6776 0.4382
0.7350 0.6810 0.4368
0.7392 0.6844 0.4352
0.7434 0.6877 0.4338
0.7476 0.6911 0.4322
0.7518 0.6945 0.4307
0.7560 0.6979 0.4290
0.7602 0.7013 0.4273
0.7644 0.7047 0.4258
0.7686 0.7081 0.4241
0.7729 0.7115 0.4223
0.7771 0.7150 0.4205
0.7814 0.7184 0.4188
0.7856 0.7218 0.4168
0.7899 0.7253 0.4150
0.7942 0.7288 0.4129
0.7985 0.7322 0.4111
0.8027 0.7357 0.4090
0.8070 0.7392 0.4070
0.8114 0.7427 0.4049
0.8157 0.7462 0.4028
0.8200 0.7497 0.4007
0.8243 0.7532 0.3984
0.8287 0.7568 0.3961
0.8330 0.7603 0.3938
0.8374 0.7639 0.3915
0.8417 0.7674 0.3892
0.8461 0.7710 0.3869
0.8505 0.7745 0.3843
0.8548 0.7781 0.3818
0.8592 0.7817 0.3793
0.8636 0.7853 0.3766
0.8681 0.7889 0.3739
0.8725 0.7926 0.3712
0.8769 0.7962 0.3684
0.8813 0.7998 0.3657
0.8858 0.8035 0.3627
0.8902 0.8071 0.3599
0.8947 0.8108 0.3569
0.8992 0.8145 0.3538
0.9037 0.8182 0.3507
0.9082 0.8219 0.3474
0.9127 0.8256 0.3442
0.9172 0.8293 0.3409
0.9217 0.8330 0.3374
0.9262 0.8367 0.3340
0.9308 0.8405 0.3306
0.9353 0.8442 0.3268
0.9399 0.8480 0.3232
0.9444 0.8518 0.3195
0.9490 0.8556 0.3155
0.9536 0.8593 0.3116
0.9582 0.8632 0.3076
0.9628 0.8670 0.3034
0.9674 0.8708 0.2990
0.9721 0.8746 0.2947
0.9767 0.8785 0.2901
0.9814 0.8823 0.2856
0.9860 0.8862 0.2807
0.9907 0.8901 0.2759
0.9954 0.8940 0.2708
1.0000 0.8979 0.2655
1.0000 0.9018 0.2600
1.0000 0.9057 0.2593
1.0000 0.9094 0.2634
1.0000 0.9131 0.2680
1.0000 0.9169 0.2731];
end

function plot_hinge_activation_figure(c, theta)

gfrac_vals = [0, 0.2, 0.4, 0.6, 0.8];
figPos = [200 300 900 350];

% Domains
u = linspace(-0.1, 0.1, 1000);
R = linspace(0, 1, 1000);

% Smooth hinge
D = @(x) 0.5 .* (x + sqrt(x.^2 + c));

% Input function
uR = (theta - R) ./ theta;

% Panel A data
hard_hinge = max(u, 0);
smooth_hinge = D(u);

% Panel B data
GAMMA = zeros(numel(gfrac_vals), numel(R));
for i = 1:numel(gfrac_vals)
    gfrac = gfrac_vals(i);
    GAMMA(i,:) = D(uR) .* (1 - gfrac);
end

% Custom colors (normalized RGB)
cols = [
    27,158,119;
    217,95,2;
    117,112,179;
    231,41,138;
    166,118,29
] / 255;

smooth_col = [217,95,2] / 255; % for Panel A

% Figure
figure('Color','w', 'Position', figPos);
ax = gobjects(2,1);

% ---------------- Panel A ----------------
ax(1) = subplot(1,2,1); hold(ax(1), 'on');

plot(u, hard_hinge, '-', 'LineWidth', 1.5, 'Color', 'k');
plot(u, smooth_hinge, '-', 'LineWidth', 2.5, 'Color', smooth_col);

set(gca,'FontSize',12,'Box','on','LineWidth',1);

xlabel('signal, $u$','Interpreter','latex','FontSize',18);
ylabel('activation signal, $D(u)$','Interpreter','latex','FontSize',20);

legend({'hard hinge', 'smooth hinge approx'}, ...
    'FontSize',10,'Location','northwest','Box','on');

% ---------------- Panel B ----------------
ax(2) = subplot(1,2,2); hold(ax(2), 'on');

for i = 1:numel(gfrac_vals)
    plot(R, GAMMA(i,:), ...
        'LineWidth', 2, ...
        'Color', cols(i,:));
end

set(gca,'FontSize',12,'Box','on','LineWidth',1);

xlabel('activation-normalised regulatory factor, $R/\tilde{R}$','Interpreter','latex','FontSize',18);
ylabel({'guard activation factor,', ...
    '$\Gamma(G,R)$'}, ...
    'Interpreter','latex','FontSize',20);
ylim([0,1])
legendStrings = arrayfun(@(x) ...
    ['$G/G_{\max} = ' num2str(x,'%.2g') '$'], ...
    gfrac_vals, 'UniformOutput', false);

legend(legendStrings, ...
    'Interpreter','latex', ...
    'FontSize',10, ...
    'Location','northeast', ...
    'Box','on');

% Panel labels
labels = {'A','B'};
for i = 1:2
    axes(ax(i)); %#ok<LAXES>
    text(0, 1.01, labels{i}, 'Units','normalized', ...
        'FontWeight','bold', 'FontSize',16, ...
        'HorizontalAlignment','left', 'VerticalAlignment','bottom');
end

end
```
