## Supplementary material for "Guarding versus self-guarding in innate immunity": Fig. S1

### Guarding versus self-guarding in innate immunity: Supplementary material

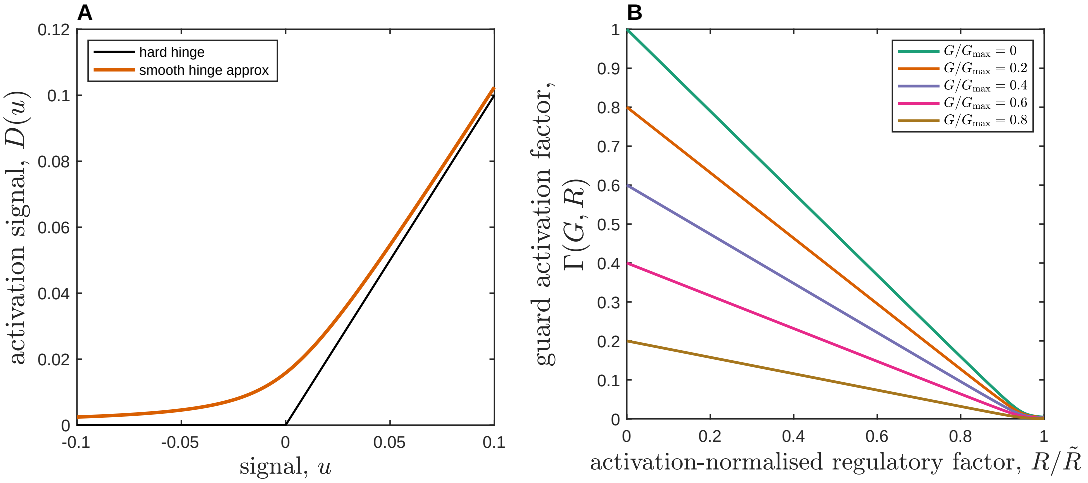


Figure 1: Illustration of the smooth hinge function and guard activation dynamics described in the Methods. (A) The smooth approximation to the hinge function, $D(u)$, used in the model (orange), compared to the hard hinge $\max(u, 0)$ (black). The smoothing parameter $c = 10^{-3}$ ensures a continuous transition around $u = 0$, avoiding a sharp activation threshold. (B) The guard activation factor $\Gamma(G, R) = D(u(R))\left(1 - \frac{G}{G_{\max}}\right)$, shown as a function of the activation-normalised regulatory factor $R/\tilde{R}$ for different fixed values of $G/G_{\max}$. Activation increases as $R/\tilde{R}$ declines below one, reflecting increasing damage, while larger values of $G/G_{\max}$ reduce the activation signal through the saturating factor $1 - G/G_{\max}$.
